# PI31 is a positive regulator of 20S immunoproteasome assembly

**DOI:** 10.1101/2025.01.15.633194

**Authors:** Jason Wang, Abbey Kjellgren, George N. DeMartino

**Affiliations:** Department of Physiology, University of Texas Southwestern Medical Center, 5323 Harry Hines Boulevard, Dallas, TX 75390-9040

## Abstract

PI31 (<u>P</u>roteasome Inhibitor of <u>31</u>,000 Da) is a 20S proteasome-binding protein originally identified as an inhibitor of *in vitro* 20S proteasome activity. Although recent studies have provided a detailed structural basis for this activity, the physiologic significance of PI31-mediated proteasome inhibition remains uncertain and alternative cellular roles for PI31 have been described. Here we report a role for PI31 as a positive regulator for the assembly of the 20S immuno-proteasome (20Si), a compositionally and functionally distinct isoform of the proteasome that is poorly inhibited by PI31. Genetic ablation of PI31 in mammalian cells had no effect on the cellular content or activity of constitutively expressed proteasomes but reduced the cellular content and activity of interferon-γ-induced immuno-proteasomes. This selective effect is a consequence of defective 20Si assembly, as indicated by the accumulation of 20Si assembly intermediates. Our results highlight a distinction in the assembly pathways of constitutive and immuno-proteasomes and indicate that PI31 plays a chaperone-like role for the selective assembly of 20S immunoproteasomes.

## INTRODUCTION

The proteasome is responsible for the degradation of most intracellular proteins in eukaryotes ^1^. Accordingly, proteasome-dependent protein degradation mediates or regulates nearly every aspect of normal cellular physiology by controlling levels of proteins responsible for given processes. Despite the structural diversity of proteasome substrates and the breadth of different physiologic states under which they are degraded, proteasome function is highly selective and tightly regulated ^2^. This selectivity and regulation is exerted by multiple mechanisms, including regulated ubiquitylation of substrates that promotes selective proteasome-substrate interaction ^3,4^, regulated posttranslational modifications of the proteasome that directly affect its catalytic activity ^5–7^, and regulation of cellular proteasome content in response to changing physiologic conditions ^8,9^. An under-appreciated mode of proteasome regulation is the structural diversity of the proteasome itself ^10^. The proteasome is a modular protease system consisting of several variants of a protease complex (20S proteasome or Core Particle) and multiple regulatory proteins to which they bind (e.g., 19S/PA700, PA28αβ, PA28γ, PA200, p97 and PI31) ^11–17^. The resulting holoenzymes feature distinct functional and regulatory properties imparted by their unique compositions. For example, the 20S-19S/PA700 holoenzyme (commonly referred to as the 26S proteasome) degrades ubiquitylated proteins by a mechanism linked to ATP hydrolysis ^2,18^. In contrast, 20S-PA28 complexes degrade certain non-ubiquitylated proteins in an ATP-independent fashion ^19,20^. Additional structural and functional proteasome diversity is conferred by holoenzymes containing copies of two different regulators ^21,22^.

20S proteasomes are the common element of all proteasome holoenzymes. The 20S proteasome is a cylinder-shaped particle composed of four axially stacked hetero-heptameric rings ^23,24^. Each of the two identical outer rings is composed of seven different but homologous α-type subunits while each of the two identical inner rings is composed of seven different but homologous β-type subunits ^23,25^. Three of the seven β subunits (β1c, β2c and β5c) feature N-terminal threonine residues that function as catalytic nucleophiles with varying specificities for peptide bond hydrolysis (cleavage after acidic, basic/neutral, and small hydrophobic residues, respectively) ^26^. The 20S proteasome exists in multiple isoforms that are characterized by unique complements of genetically distinct catalytic β subunits. For example, cells specialized for immune function constitutively express three so-called βi subunits (β1i, β2i and β5i), two of which (β1i and β5i) feature different cleavage specificities than their constitutive counterparts (cleavage after small hydrophobic and bulky hydrophobic residues, respectively) ^27–31^. βi subunits are also conditionally expressed in cells exposed to certain cytokines, such as interferon-γ. These interferon-induced subunits are selectively incorporated into newly synthesized proteasomes in favor of their constitutively expressed β1c, β2c and β5c counterparts ^30,32^. A specialized 20S proteasome composed of β1i/β2i subunits and β5t, a unique β5 subunit, is expressed in thymocytes ^33,34^. Other 20S proteasomes appear to have mixed populations of βc and βi catalytic subunits, although but the basis for their formation is unclear ^35^. Different 20S isoforms have different physiologic functions. For example, the catalytic specificities of immunoproteasomes promote the production of antigenic peptides required for immune surveillance ^36^ and may be required for the clearance of oxidatively damaged proteins ^37,38^. Likewise, the β5t subunit of thymo-proteasomes promotes the positive selection of CD8^+^ T cells ^33,34^. Thus, different 20S isoforms significantly expands the functional complexity of the proteasome system.

The 20Sc proteasome is assembled from its 28 component subunits by a highly ordered, stepwise process that is aided by five dedicated chaperones, as well as by chaperone-like functions of N-terminal pro-peptides of catalytic β subunits ^39–42^. A current model of 20S biogenesis, derived from extensive genetic, biochemical and structural data, indicates that the process begins with the formation of a complete α1-α7 subunit ring. This process is mediated by two heterodimeric chaperones, PAC1/PAC2 and PAC3/PAC4 (Pba1/Pba2 and Pba3/Pba4, respectively in yeast) that remain bound to opposite faces of the completed α-subunit ring^43^. The chaperone POMP (Ump1 in yeast) then binds to the PAC3/PAC4 side of the α ring and initiates β ring assembly by recruiting the pro-β2c subunit. The pro-peptide of β2c aids recruitment of the β3 subunit, which displaces PAC3/PAC4 and ejects it from the complex ^41^. The remaining β subunits are then added in rank order (β4, β5, β6, β1, and β7) in a process directed by POMP and the pro-peptide of β5c. The resulting complex, often referred to as “15S”, consists of a half-proteasome (i.e., one α-subunit ring and one β-subunit ring) featuring bound PAC1/PAC2 and POMP chaperones and catalytic β subunits with unprocessed pro-peptides. Two half-proteasome complexes then join along abutting β subunit rings to form the preholo-20S proteasome that initially retains all components of the individual 15S complexes. The fusion of two half-proteasomes induces a series of conformational changes that promotes the processing of β-subunit pro-peptides, degradation of POMP, and dissociation of PAC1/PAC2 to generate the mature 20S proteasome ^41^.

PI31 (<u>P</u>roteasome Inhibitor of <u>31</u>,000 daltons) was originally identified as an *in vitro* inhibitor of 20S proteasome activity ^44,45^. Several recent reports have defined the detailed structural basis for this function; PI31’s natively unstructured C-terminus enters the central catalytic chamber of the proteasome and directly interacts with the catalytic threonine residue of each β subunit ^46–48^. These interactions occur in a manner predicted to block hydrolytic activity of each catalytic site and are oriented in a manner that renders PI31 itself resistant to hydrolysis. The inhibitory features of PI31 have been exploited to develop small molecule, subunit-specific proteasome inhibitors based on PI31’s structure ^49^. Despite these biochemical findings, there is little evidence that PI31 functions as a global proteasome inhibitor in intact cells ^50^. In contrast, PI31 has been proposed to regulate a variety of cellular processes with no apparent relationship to one another or to mechanisms involving direct inhibition of the proteasome’s hydrolytic activity ^51–57^.

Our original characterization of PI31 as a proteasome inhibitor and our subsequent cryoEM structure of the 20S-PI31 complex were conducted with 20S proteasomes harboring constitutive catalytic β subunits (20Sc) ^44,48^. We recently compared effects of PI31 on purified 20Sc and 20S immunoproteasomes (20Si) *in vitro* and showed that PI31 had significantly attenuated inhibitory activity against 20Si ^58^. Surprisingly, this diminished inhibitory activity was largely accounted for by 20Si-catalyzed cleavage of PI31 at residues normally involved in proteasome inhibition. These findings indicate that PI31 interacts differently with the β subunits of 20Sc and 20Si and raise the possibility that that PI31 may have different cellular roles for constitutive- and immuno-proteasomes. Here we describe a role of PI31 in the assembly of 20Si induced by interferon-γ. Our data indicate that PI31 is an important factor that drives the assembly and maturation of IFNγ-induced 20S immunoproteasome.

## RESULTS

### PI31 knockout cells have normal proteasome content and activity under standard culture conditions

To investigate physiologic roles of the proteasome regulator, PI31, we ablated PI31 expression in HAP1 and HepG2 cells using CRISPR/Cas9 methodology. Disruption of the PI31 gene (PSMF1) was confirmed by sequencing (Supplemental Figure 1) and no PI31 protein was detectable by western blotting in either PI31 knockout (KO) cell line (Figure 1A). PI31 KO cells grew at indistinguishable rates from their parental counterparts under standard cell culture conditions replete with serum and nutrients (Supplemental Figure 2). Under these same conditions, the absence of PI31 had no appreciable effect on several measures of proteasome function including steady state levels of polyubiquitylated cellular proteins, rates of hydrolysis of model peptide substrates, and labeling of the catalytic β subunits with the proteasome activity probe Me_4_BodipyFL-Ahx_3_LeuVS (Figures 1B, 1C and 1D). PI31 WT and KO cells had similar levels of representative proteasome subunits (Figure 1A) and similar levels, complements, and activities of intact proteasome complexes, as judged by native polyacrylamide gel electrophoresis coupled with zymography (for activity) or western blotting (for protein levels) (Figure 1E). These results with PI31 KO cells are in accord with those of previous studies showing that depletion of PI31 content by RNAi had no appreciable effect on global cellular proteasome content or activity under normal cell culture conditions ^50^.

**Figure 1.**
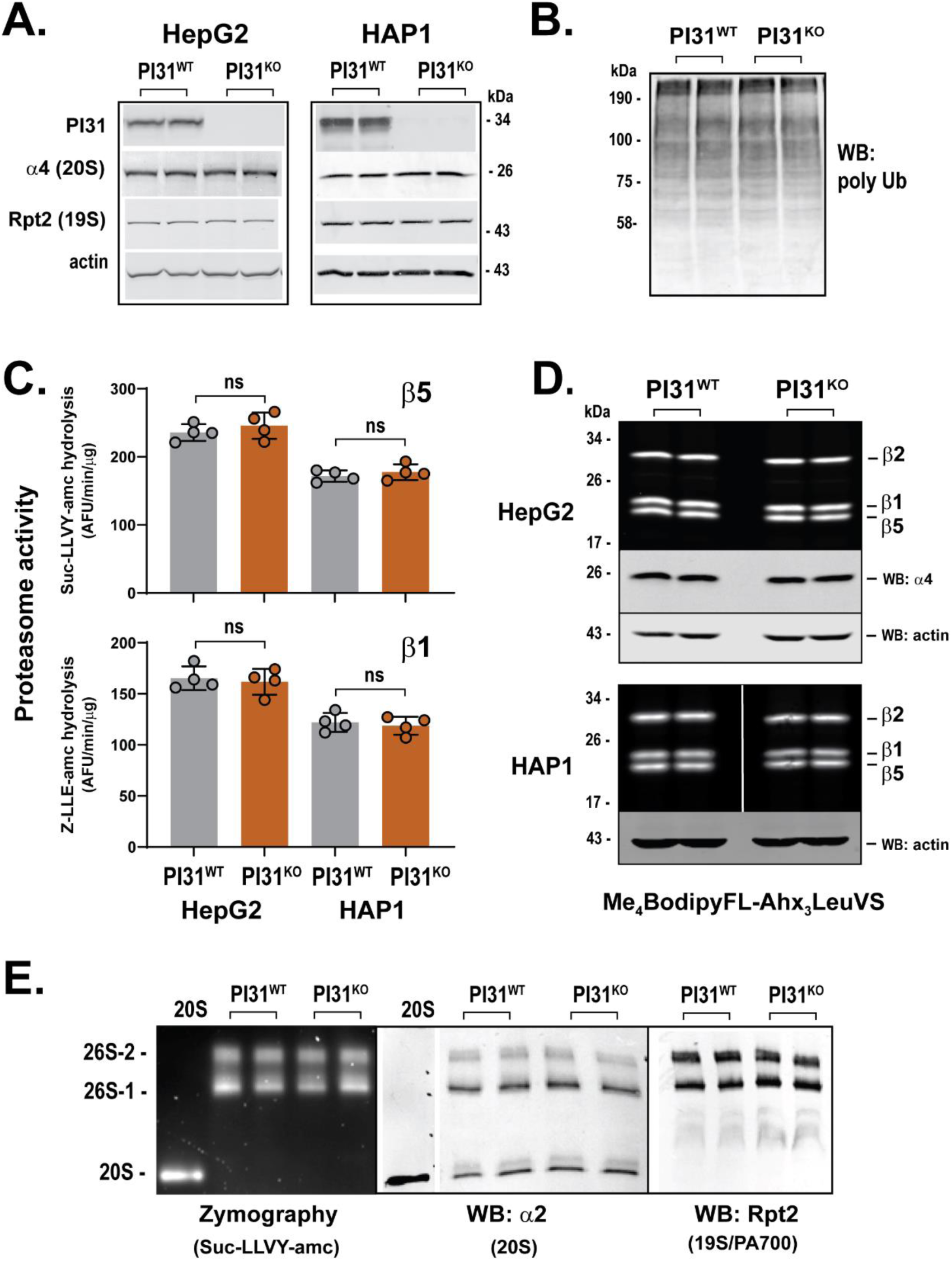
PI31 knockout cells have normal levels and activity of constitutive proteasomes. PI31 wild-type (WT) and knockout (KO) HAP1 and HepG2 cells were assessed for various features of proteasome content and activity. In all cases, extracts from WT and KO cells compare equal levels of total extract protein. **(Panel A),** Extracts of HAP1 and HepG2 PI31 WT and PI31 KO cells were subjected to SDS-PAGE and western blotting for the indicated proteins. Duplicate lanes are samples from independent biologic preparations. **(Panel B),** Extracts of PI31 WT and PI31 KO HAP1 cells were subjected to SDS-PAGE and western blotting for polyubiquitylated proteins. Duplicate lanes represent independent biologic preparations. **(Panel C),** Proteasome activity was measured using indicated substrates for extracts of PI31 WT and PI31 KO HAP1 and HepG2 cells. Each individual data point (O) represents an independent biologic experiment and is a mean value triplicate assays. Bars represent mean values ± s.d. of independent biologic experiments and were analyzed by ANOVA (*ns*: p > 0.05). **(Panel D),** Extracts of PI31 WT and PI31 KO HAP1 and HepG2 cells were treated with Me_4_BodipyFL-Ahx_3_Leu_3_VS and analyzed as described under Materials and Methods. Indicated samples were sujected to western blotting for 20S α4 subunit and actin. Duplicate lanes are samples from independent biologic experiments. **(Panel E),** Extracts of PI31 WT and PI31 KO HAP1 cells were subjected to native PAGE and analyzed by zymography using Suc-LLVY-amc substrate, or by western blotting for indicated proteins. “20S” indicates purified 20S proteasome. Duplicate lanes are samples from independent biologic experiments. The lane showing purified 20S proteasome was from the same gel and membrane as extract samples but was spliced from another part of membrane.

### PI31 knockout cells have attenuated increases of immuno-proteasome content and activity in response to interferon-γ treatment

HAP1and HepG2 cells normally express β1c, β2c, and β5c isoforms of catalytic proteasome subunits almost exclusively, and therefore contain predominately constitutive forms of functional proteasome complexes. These cells, however, up-regulate transcription of genes for β1i, β2i, and β5i subunits when exposed to interferon-γ (IFNγ). The IFNγ-induced βi catalytic subunits are selectively incorporated into newly synthesized proteasomes, commonly termed “immuno-proteasomes” (20Si), in lieu of their βc counterparts. Because we recently showed that purified PI31 interacts differently with purified 20Sc and 20Si proteasomes *in vitro* ^58^, we sought to determine if PI31 exerts different effects on constitutive- and immuno-proteasomes in intact cells. Accordingly, we treated PI31 wild-type and KO HAP1 and HepG2 cells with IFNγ and then compared their constitutive and immuno-proteasome content and activity. In the absence of IFNγ, the levels of catalytic βc subunits were indistinguishable between PI31 wild-type and KO cells (Figure 2). These results are consistent with other comparisons of proteasome content and activity between PI31 wild-type and KO cells (Figure 1). IFNγ had no general effect on the levels of βc subunits in either PI31 WT or KO cells, although protein levels declined significantly in a few instances (e.g., β1c and β5c in HAP1 WT cells) and there was an overall trend for reduction among others (Figure 2). These reductions in βc protein levels are likely a consequence of subunit turnover and the noted distinctions in these effects may reflect different clearance rates of individual βc subunits between cells during the IFNγ treatment period.

**Figure 2.**
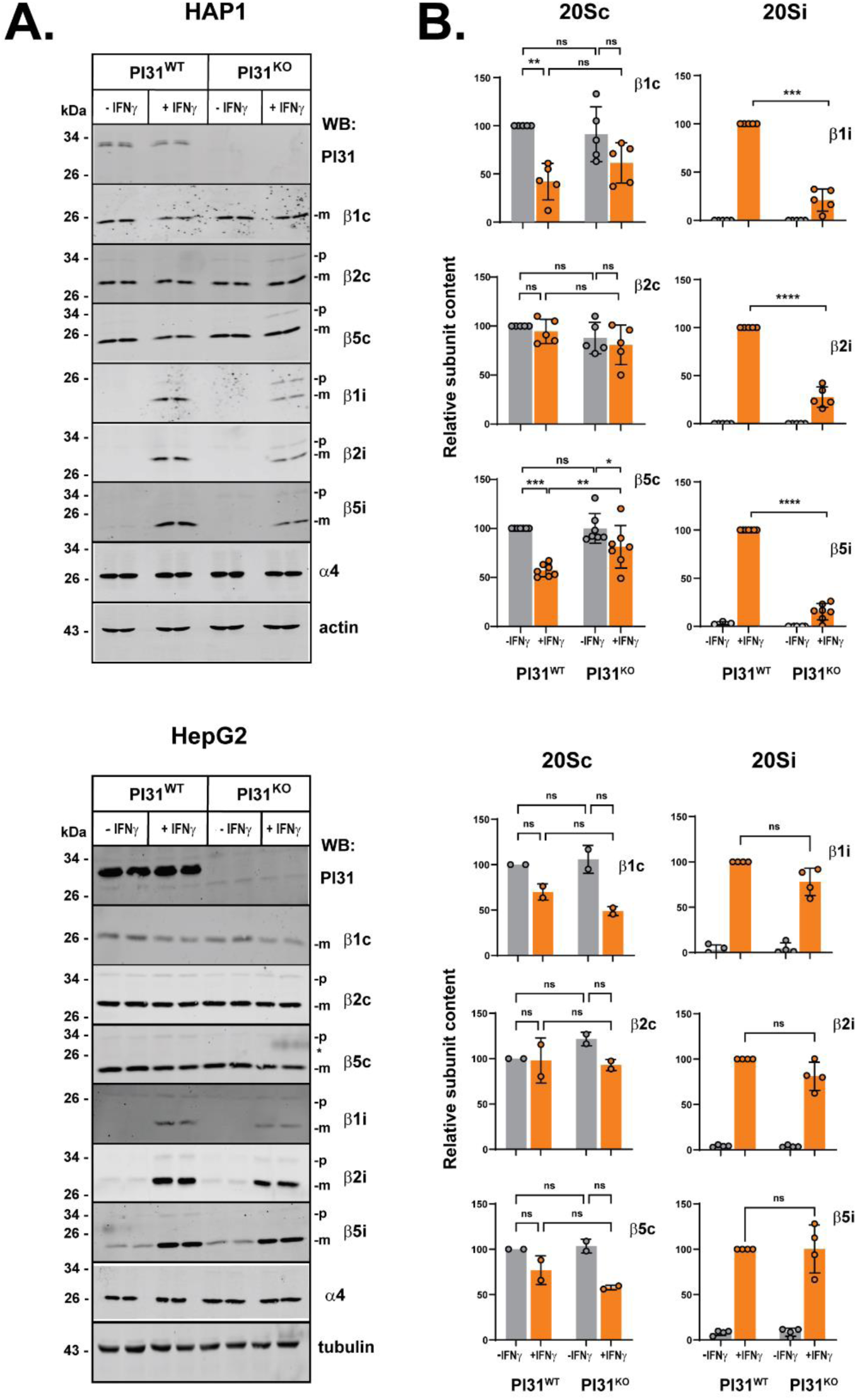
PI31 knock-out cells have reduced levels of IFNγ-induced immuno-proteasome subunits. PI31 WT and KO HAP1 (upper) and HepG2 cells (lower) were exposed to 100 U/ml human IFNγ for 24 hours. **(Panel A),** Cell extracts from treated and untreated cells were normalized for total protein and subjected to SDS-PAGE and western blotting for indicated proteins. Duplicate lanes are samples from independent biologic experiments. Bands denoted “p” and “m” represent the pro-peptide and mature forms of β subunits, respectively. **(Panel B),** Western blots of indicated 20Sc and 20Si subunits were quantified using ImageStudio (LiCOR) software from individual biologic experiments, represented by data points (O); bars represent mean values ± s.d. of independent biologic experiments; **(Panel B, left),** Subunit levels of untreated PI31 WT cells were assigned a value of 100 and subunit levels for other conditions are expressed relative to that value; **(Panel B, right),** Subunits levels of IFNγ-treated PI31 WT cells were assigned a value of 100 and subunit levels for other conditions are expressed relative to that value. Differences were analyzed by repeated measures 2-way ANOVA and Tukey’s HSD posthoc test (* p<0.05; ** p<0.01; *** p<0.001; **** p<0.0001).

As expected, IFNγ treatment for as little as 24 hours greatly increased expression of each βi subunit in both PI31 wild-type HAP1 and HepG2 cells (Figure 2). IFNγ also increased βi content in PI31 KO cells, but the degree of increase was attenuated compared to that in wild-type cells. Attenuation was especially prominent in HAP1 cells and was observed for each βi subunit (Figure 2). PI31 KO HepG2 cells also featured a diminished βi subunit content in response to IFNγ, but the magnitude of the effect was smaller and more variable than in HAP1 cells and did not reach statistical significance. Despite the distinction of this latter effect between the HAP1 and HepG2 cells, further analysis described below shows that these PI31 KO cells share similar defects in the assembly of βi subunits into mature immunoproteasomes.

To determine functional consequences of the lack of PI31 on content and activity of intact immunoproteasomes, we measured and compared proteasome activity in extracts of PI31 wild-type and KO cells that had or had not been treated with IFNγ. IFNγ treatment had little or no effect on rates of hydrolysis of Suc-LLVY-amc, a substrate cleaved by both the β5c and β5i subunits of respective constitutive and immuno-proteasomes, in either PI31 wild-type or KO cells (Figure 3). Statistically significant increases were noted in some instances, but the magnitude of these effects was modest. In contrast, IFNγ treatment of PI31 wild-type cells promoted a large increase (2-6 fold) in the hydrolysis of Ac-ANW-amc and Ac-PAL-amc, peptides cleaved with high specificity by the β5i and β1i subunits, respectively, of immuno-proteasomes (Figure 3 and Supplemental Figure 3). These IFNγ-induced immunoproteasome activities, however, were significantly attenuated (40-60% reductions) in PI31 KO HAP1 and HepG2 cells (Figure 3). Thus, genetic ablation of PI31 expression reduced IFNγ-induced immunoproteasome activity in both HAP1 and HepG2 cells, despite its dissimilar effect on their overall levels of βi subunit content. This result suggests that the subunit content, *per se*, is not an accurate reflection of intact, functional proteasome complexes. To confirm the role of PI31 on IFNγ-induced immunoproteasome activity, we employed RNAi as an independent method of reducing cellular PI31 content. HepG2 cells were transfected with siRNAs targeting PI31 prior to treatment with IFNγ. RNAi reduced PI31 content to undetectable levels (Supplemental Figure 4). Consistent with results in PI31 KO cells and with previously reported data with HEK293 cells, RNAi of PI31 had little effect on the content or activity of constitutive proteasomes in the absence of IFNγ ^50^. In contrast, IFNγ-induced immunoproteasome activity was significantly reduced in PI31 knockdown cells (Supplemental Figure 4). This effect was accompanied by a reduction in the content of βi subunits. These results show that orthogonal methods of reducing cellular PI31levels produce similar results on inhibition of immuno-proteasome activity.

**Figure 3.**
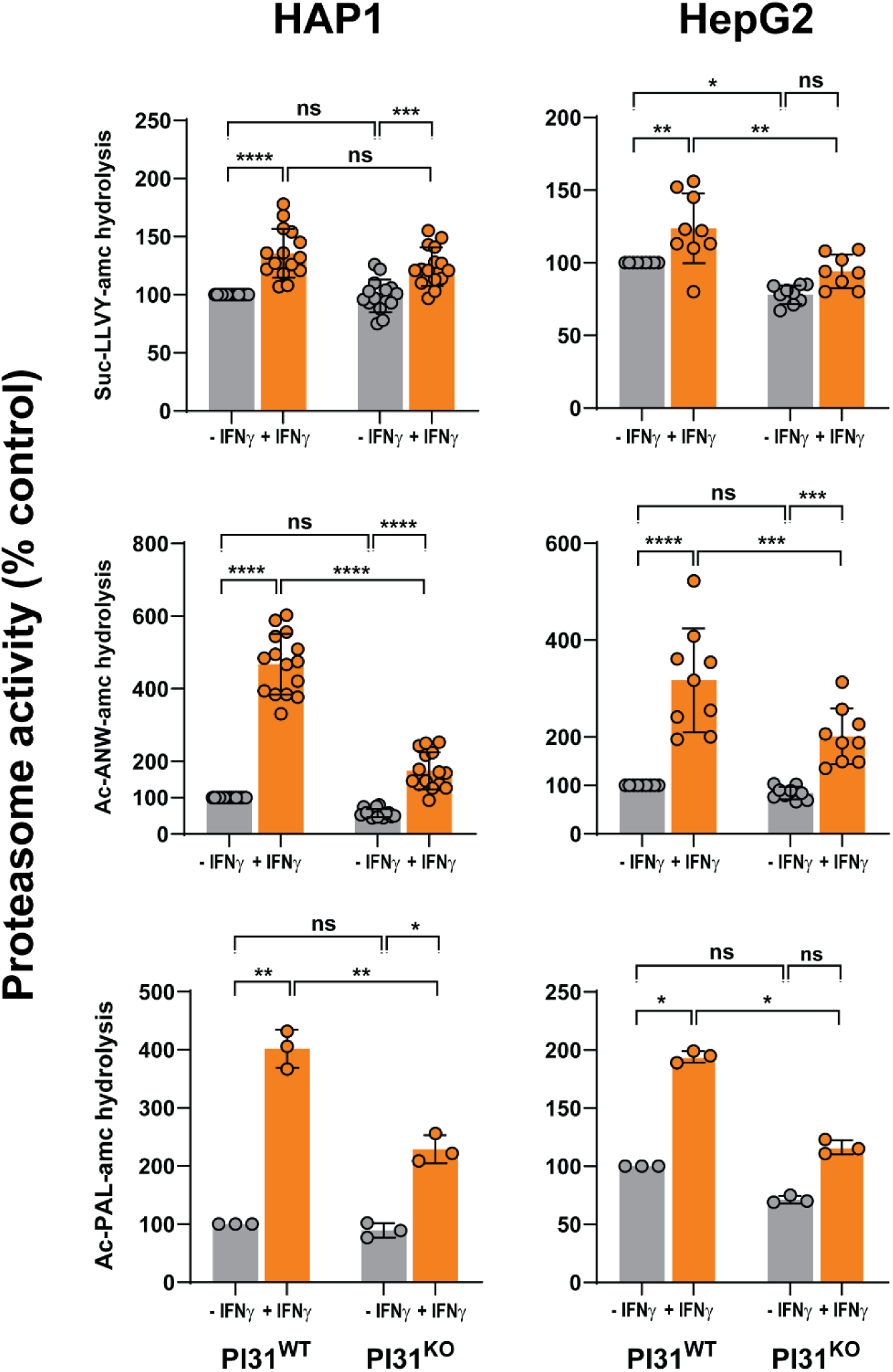
PI31 knockout cells have reduced IFNγ-induced immuno-proteasome activity. PI31 WT and PI31 KO HAP1 and HepG2 cells were treated with 100 U/ml human IFNγ for 24 hours. Cell extracts were normalized for total protein and assayed for proteasome activity using Suc-LLVY-amc, Ac-ANW-amc, and Ac-PAL-amc substrates, as described under Materials and Methods. Mean activity values for extracts from untreated wild-type cells were measured and assigned a value of 100. Activities of other conditions are expressed relative to that value. Activities of individual biologic preparations (O) are mean values of triplicate assays. Bars represent mean values ± s.d. of independent biological experiments. Differences were analyzed by repeated measures 2-way ANOVA and Tukey’s HSD posthoc test (* p<0.05; ** p<0.01; *** p<0.001; **** p<0.0001).

To gain deeper insight into the basis of the diminished IFNγ-induced immuno-proteasome activity in PI31 KO cells, we subjected extracts of IFNγ-treated and IFNγ-untreated PI31 wild-type and KO cells to native PAGE and analyzed the gels by zymography (for proteasome activity) and western blotting (for the content and composition of proteasome complexes). Most zymographic activity in untreated wild-type and knockout cells was accounted for by constitutive 26S proteasomes (Figure 4A, lanes 1, 3, 31, 33). Consistent with analysis of activities measured in whole cell extracts, zymography of IFNγ-treated wild-type cells displayed substantially increased activities against immunoproteasome-selective substrates such as Ac-ANW-amc (selective for the β5i subunit; compare lanes 5 and 6; 36 and 37) and Ac-PAL-amc (selective for the β1i subunit; compare lanes 41 and 42). These immunoproteasome activities were accounted for by both 26S proteasomes and a band (Band 1) that migrated slightly slower than pure 20S proteasome (Figure 4A; compare lanes 35 and 36-37, and lanes 40 and 41-41). Western blotting with an antibody that recognizes multiple α and β subunits of 20S proteasome revealed that the area of the gel near Band 1 activity contains several closely migrating bands (Figure 4A, lanes 13-17, and lanes 45-49) that likely represent a mixture of immature preholo-20S, catalytically latent mature 20S, and 20S-PA28αβ holoenzyme complexes. Identification of these proteasome forms as components of Band 1 is supported by western blotting with antibodies against PA28αβ and PAC1 (Figure 4A, lanes 9-12, 22-25, and 54-57). Additional evidence for the designations of these complexes is presented below. Notably, the activity of the 20S-PA28αβ complex was increased by IFNγ to a proportionally greater extent than that of 26S proteasomes and therefore appears to account for most of the IFNγ-induced increase of immuno-proteasome activity. A selective increase in 20Si-PA28 activity in response to interferon-γ treatment has been reported previously ^59^. Although the expected increase of PA28αβ expression by interferon-γ observed in these experiments (Figure 4A and Supplemental Figure 5) may play a role in the preferential increase of 20Si-PA28αβ content and activity, PA28αβ levels *per se* do not appear to limit formation of this holoenzyme because PA28 unassociated with the proteasome was observed in both treated and untreated cells (Figure 4A, lanes 9 and 10). Nevertheless, IFNγ-induced zymographic immuno-proteasome activities were markedly attenuated in both PI31 KO HAP1 and HepG2 cells (Figure 4). These results are consistent with the attenuated IFNγ-induced immuno-proteasome activity of PI31 KO cells measured in whole cell extracts and suggest that PI31 is required for normal induction of immuno-proteasome activity in response to interferon-γ (Figure 4A, compare lanes 6 and 8, 37 and 39, 42 and 44). PI31 levels were not affected by IFNγ treatment indicating that the basal content of PI31 was sufficient to support this proposed role (Supplemental Figure 5).

**Figure 4.**
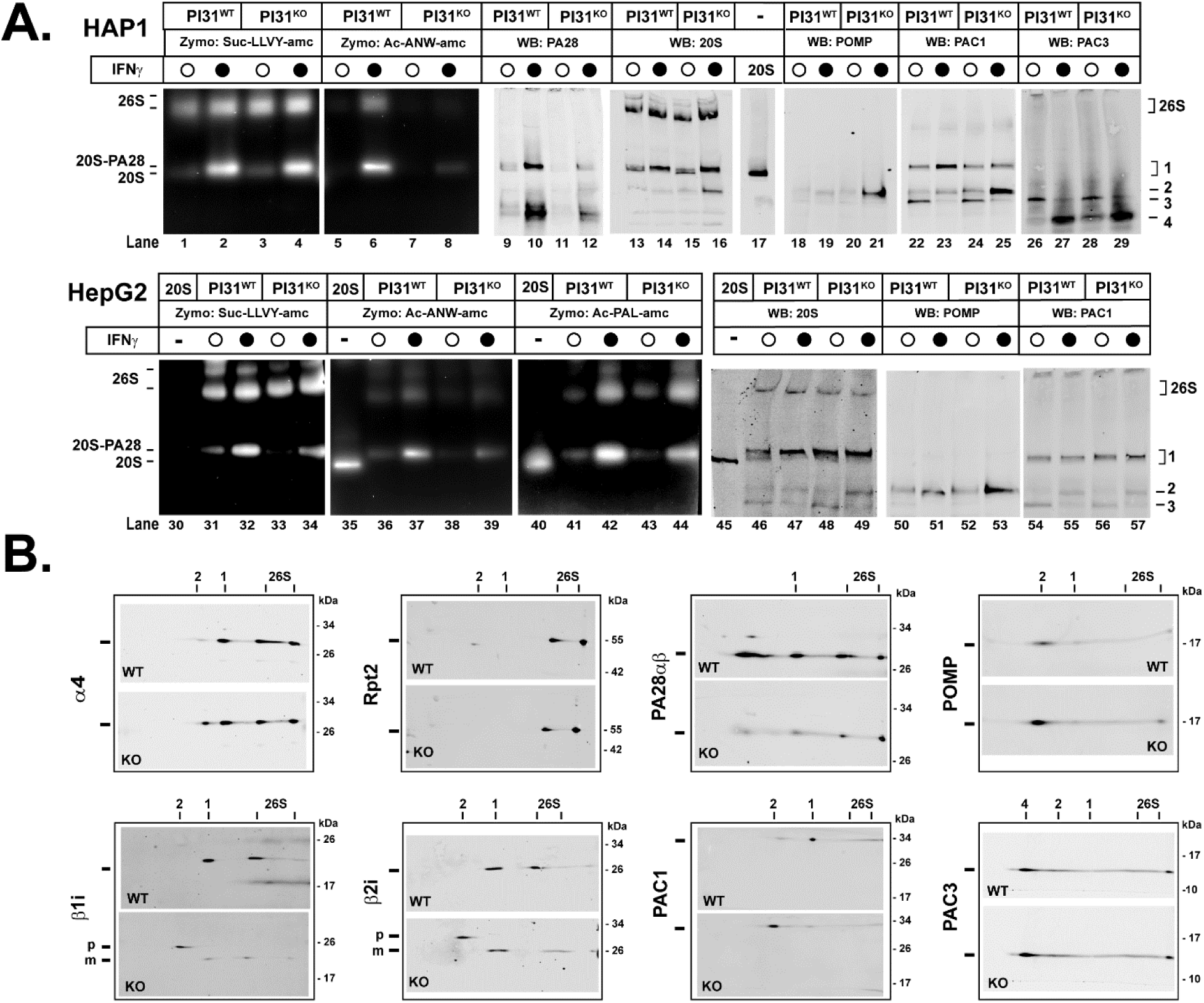
PI31 knockout cells have altered content and activity of IFNγ-induced proteasome complexes. PI31 WT and PI31 KO HAP1 and HepG2 cells were exposed to 100 U/ml human IFNγ (λ) or control buffer (O) for 24 hours. **(Panel A),** Cell extracts were normalized for total protein and subjected to native PAGE for zymography with indicated proteasome substrates or western blotting for indicated proteins. Purified 20S proteasome was electrophoresed as a standard. The lane showing purified 20S proteasome was from the same gel and membrane as extract samples but was spliced from another part of membrane. Complexes denoted 26S and 1-4, are described in the text. **(Panel B),** Extracts of PI31 WT and KO HAP1 cells treated with interferon-γ and subjected to native PAGE as in Panel A, followed by second dimension SDS-PAGE and western blotting for indicated proteins. Positions denoted 26S, 1, and 2 correspond to those of the first dimension native gel. “p-” and “m” indicate positions of the unprocessed pro-peptide and mature forms of indicated βi subunits. Similar results were obtained in four independent experiments.

In contrast to zymography, which allowed for identification of immuno-proteasome-selective activity, we were unable to directly distinguish constitutive and immuno-proteasome content on native gels because immuno-proteasome-specific antibodies available to us reacted poorly with intact native complexes. To circumvent this technical limitation, we subjected proteasome complexes resolved on native gels to second dimension SDS-PAGE and western blotting. This method confirmed identities assigned to these complexes on native gels and specifically identified immuno-proteasomes. For example, Rpt2, a subunit of the 19S/PA700 regulatory complex common to both constitutive and immuno-26S proteasomes, was identified at horizontal positions of the SDS-gel that aligned with complexes identified as 26S proteasomes on native gels. Likewise, α4, a subunit common to both constitutive and immuno-20S and 26S proteasomes, was detected at horizontal positions of the SDS gel that aligned with native gel bands assigned those identities. Consistent with the analysis of native gels alone, PA28αβ and 20S were detected at coincident horizontal positions on SDS-gels that in turn aligned with the IFNγ-induced activity band from the native gel. This finding supports the conclusion that the activity of Band 1 is accounted for by a 20S-PA28 holoenzyme. PA28 was also identified in the regions corresponding to 26S proteasomes, suggesting the presence of 20S-19S/PA700-PA28 hybrid complexes in these bands of zymographic activity. PA28 was also identified in the position characteristic of a proteasome-unbound form. Most importantly, mature βi subunits were identified in horizontal positions that aligned with those of both 20S/20S-PA28 (Band 1) and 26S proteasome complexes. Notably, the βi content in these bands was lower in IFNγ-treated PI31 KO cells than in corresponding wild-type cells. Collectively, these results support the conclusion that the reduced IFNγ-stimulated immuno-proteasome activity of PI31 KO cells is a consequence of decreased immuno-proteasome content.

### Cells lacking PI31 accumulate 20S immuno-proteasome assembly intermediates containing 20S assembly chaperones and unprocessed βi subunits in response to interferon-γ

In addition to detecting fully-assembled, catalytically competent proteasomes, native PAGE also can identify intermediate complexes of the 20S assembly process. The native PAGE analysis of proteasome complexes described above revealed several bands that reacted with antibodies against 20S proteasome and migrated more rapidly than the purified 20S proteasome standard. We reasoned that these bands likely represent such 20S assembly intermediates (Figure 4A, bands denoted 2, 3 and 4). For example, one of these bands (denoted Band 3, Figure 4A, lanes 22, 24, 26, and 28) featured both PAC1 (a monitor of the PAC1/PAC2 heterodimer) and PAC3 (a monitor of the PAC3/PAC4 heterodimer) and therefore likely represents an early-stage complex composed of an intact α-subunit ring. The decreased content of Band 3 and the appearance of a new PAC3-containing band (Band 4, lanes 23, 25, 27, and 29) upon IFNγ treatment, may indicate the progression of early-stage immuno-proteasome assembly. Notably, another of these bands (Band 2, Figure 4A, lanes 16 and 49) accumulated in samples from IFNγ-treated PI31KO HAP1 and HepG2 cells. This finding suggests that attenuated immuno-proteasome content and activity of IFNγ-treated PI31 KO cells may be a consequence of defective or retarded 20Si assembly. To test this possibility, we examined the composition of the putative assembly intermediate in greater detail. Western blotting of native polyacrylamide gels showed that in addition to subunits of 20S proteasome, Band 2 was detected by antibodies against the established 20S assembly chaperones POMP and PAC1 (Figure 4A, lanes 18-25 and 50-57), but lacked the 20S chaperone PAC3 (Figure 4A, lanes 26-29). Levels of the POMP and PAC1 chaperones in Band 2 were highest in extracts from IFNγ-treated PI31 KO cells (Figure 4A, lanes 21, 25, and 53). Western blotting of whole cell extracts confirmed that POMP levels were approximately 2-fold greater in IFNγ-treated PI31 KO cells than in untreated cells (Supplemental Figure 5). The protein composition of Band 2 is characteristic of the 15S half-proteasome assembly complex. In addition to the POMP and the PAC1/PAC2 chaperones, this late-stage 20S assembly intermediate is also known to contain unprocessed pro-peptide forms of catalytic β subunits. To further characterize Band 2 from native gels, we again analyzed second dimension SDS gels by western blotting. Both POMP and PAC1 were detected at a horizontal position of the SDS gel that aligned with Band 2 on the native gel. In agreement with the relative intensities of the western blotting signals in native gels, each of these proteins was present at higher levels in IFNγ-treated PI31 KO cells than in IFNγ-treated PI31 wild-type cells. Unprocessed, pro-peptide forms of βi subunits also were detected in the POMP/PAC1-containing Band 2 after second dimension SDS-PAGE. These unprocessed βi subunits featured the expected higher molecular weights than their processed mature forms that were located at positions that aligned with the fully assembled 20S (Band 1) and 26S proteasomes on native gels. Collectively, these findings support the conclusion that an assembly intermediate of IFNγ-induced 20Si proteasomes accumulates in PI31 KO cells and that this intermediate is similar or identical to the 15S half-proteasome.

Defective IFNγ-induced 20Si proteasome assembly was also demonstrated using glycerol density gradient centrifugation, an orthogonal method of identifying and comparing the content and activities of different proteasome complexes via functional assays and western blotting of gradient fractions (Figure 5). This method confirmed key findings obtained by the native PAGE analysis described above. Thus, i) untreated PI31 wild-type and KO cells had similar overall levels of constitutive proteasome activity and this activity was accounted for mainly by constitutive 26S holoenzymes (Figure 5A, upper left and upper right); ii) IFNγ greatly increased immuno-proteasome activity in PI31 wild-type cells (Figure 5A, lower left); iii) IFNγ-stimulated immuno-proteasome activity was accounted for by both 26S and 20S-PA28 complexes, but the latter complex was increased to a proportionally greater extent than the former (Figure 5A, lower left), and; iv) IFNγ-induced immuno-proteasome activity was markedly reduced in PI31 KO cells (Figure 5, lower left and lower right).

**Figure 5.**
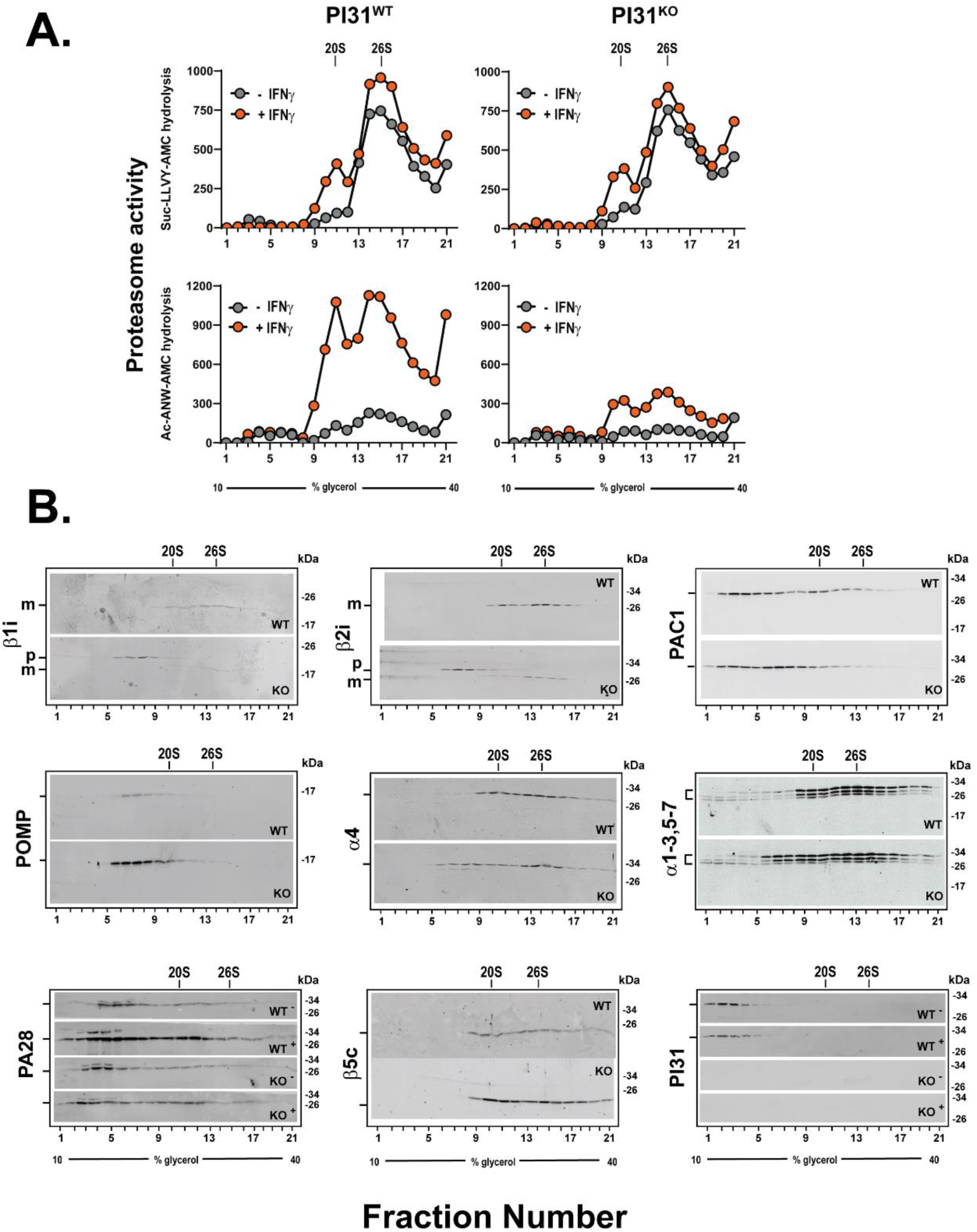
PI31 knockout cells have altered content and activity of IFNγ-induced proteasome complexes. PI31 WT and PI31 KO HAP1 were exposed to 100 U/ml human IFNγ for 24 hours. Cell extracts were normalized for total protein and subjected to 10-40% glycerol density gradient centrifugation. **(Panel A),** Gradient fractions were assessed for proteasome activity with indicated substrates. **(Panel B),** Gradient fractions from Panel A were subjected to western blotting for indicated proteins. Unless otherwise noted by “-” or “+”, blots show samples from treatments with interferon-γ. 26S, 20S-PA28, and 20S show sedimentation positions of purified complexes as standards. “p-” and “m” indicate unprocessed pro-peptide and mature forms, respectively, of indicated βi subunits. Similar results were obtained in four independent experiments for activity measurements and in three independent experiments for western blotting.

To further correlate these various proteasome activities with specific proteasome complexes we subjected gradient fractions to western blotting (Figure 5B). As expected, IFNγ-treated PI31 wild-type cells featured bimodal distribution profiles of representative 20S proteasome subunits, including α subunits and mature βc, and βi subunits. The distribution profiles of these proteins were coincident with peaks of proteasome activity attributed to 26S (Fractions 13-17) and 20S-PA28 (fractions 8-12) holoenzymes. The latter assignment was supported by a peak of PA28αβ that was coincident with both 20S subunits and proteasome activity. The level of PA28αβ in the latter peak was greater in IFNγ-treated than untreated PI31 wild-type cells, a finding consistent with the conclusion that the 20Si-PA28 holoenzyme is preferentially formed in response to interferon-γ treatment. The content and distribution of βi subunits in IFNγ-treated PI31wild-type and KO cells were appreciably different. Thus, the level of mature βi subunit peaks of 20S and 26S complexes was reduced in PI31 KO cells, and unprocessed pro-peptide forms of these subunits accumulated in slower sedimenting fractions (Figure 5B, fractions 6-8). These fractions also contained a peak of α subunits not present in IFNγ-treated wild type cells, as well as enhanced peaks of POMP and PAC1. These findings are consistent with the accumulation of a complex similar to that identified as the 15S half-proteasome (Band 2) by the native PAGE analysis described above (Figure 4). Collectively, these results are in excellent accord with those of the native PAGE analysis and support the conclusion that IFNγ-treated PI31 KO cells are defective in the assembly of 20Si proteasome.

### Knockdown of POMP reveals a distinct role for PI31 in immuno-proteasome assembly

The aberrant accumulation of a 20Si assembly intermediate in PI31 KO cells suggests that PI31 plays a normal role in 20Si assembly and maturation. To gain additional insight into this role, we compared the effects of PI31 knockout on IFNγ-induced immuno-proteasome content and activity with those resulting from depletion of POMP, an established 20S assembly chaperone. siRNAs directed against POMP decreased cellular POMP content to less than 5% of control levels in both IFNγ-treated and IFNγ-untreated PI31 wild-type and KO cells (Figure 6A). Depletion of POMP inhibited IFNγ-induced immuno-proteasome activities to a significantly greater extent than did knockout of PI31 (> 80% versus ∼ 50%, respectively), but depletion of POMP in PI31 KO cells had no significantly greater inhibitory effect than depletion of POMP alone (Figure 6B). POMP knockdown decreased the steady-state levels of all βc and βi catalytic subunits in both IFNγ-treated and IFNγ-untreated PI31 wild-type cells (Figure 6B). This result was mirrored by the effect of POMP knockdown on the hydrolysis of Suc-LLVY-amc, a measure of both constitutive and immuno-proteasome activities. Moreover, the previously described decrease of IFNγ-induced βi subunit content in PI31 KO cells was more pronounced when these cells were also depleted of POMP (Figure 6A).

**Figure 6.**
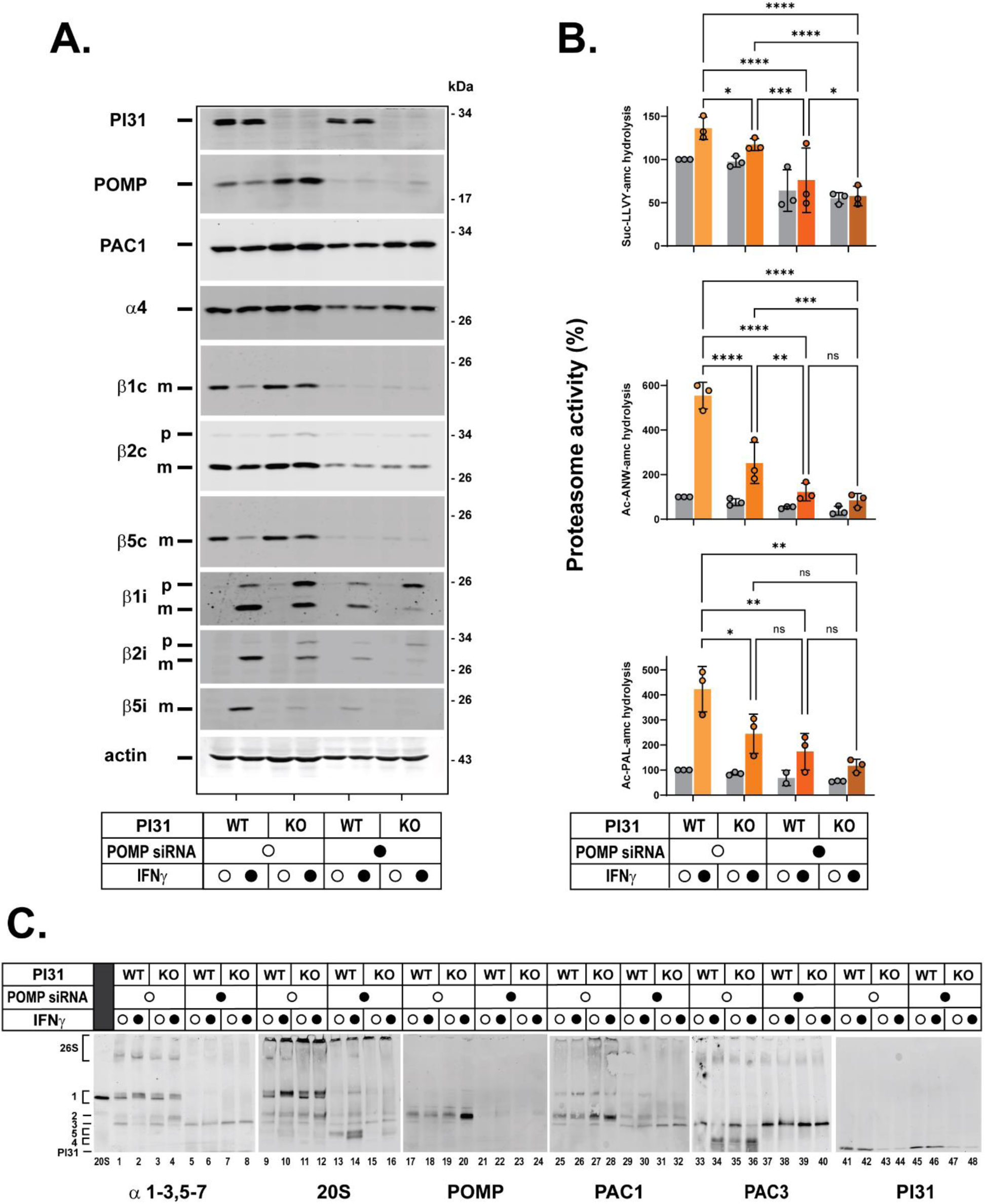
Depletion of PI31 and POMP have distinct effects on IFNγ-induced proteasome activity and content. PI31 WT and PI31 KO HAP1 cells were treated with POMP coding or non-coding siRNAs for 24 hours prior to treatment for 24 hour with 100 U/ml human IFNγ or control media. Cell extracts were normalized for total protein and subjected to the following analyses: **(Panel A),** Extracts were subjected to SDS-PAGE and western blotting for the indicated proteins; **(Panel B),** Extracts were assayed for proteasome activity against the indicated substrates. Mean activity values for extracts from untreated wild-type cells were assigned a value of 100 and activities of other conditions are expressed relative to that value. Activities of individual biologic preparations (O) are mean values of triplicate assays. Bars represent mean values ± s.d. of independent biological experiments. Differences were analyzed by repeated measures 2-way ANOVA and Tukey’s HSD posthoc test (* p<0.05; ** p<0.01; *** p<0.001; **** p<0.0001); **(Panel C),** Cell extracts were subjected to 3-8% native PAGE and western blotting for indicated proteins. Bands denoted 26S and 1-5 are described in the text and correspond to those shown in Figure 4. Similar results were obtained in six independent experiments for 20S, POMP and PI31, in three independent experiments for PAC1 and PAC3, and two independent experiments for α1-3,5-7.

To further examine the relative effects of POMP and PI31 on IFNγ-induced immuno-proteasome assembly, extracts from cellular conditions described above were electrophoresed on native gels and blotted with antibodies that detect various 20S proteasome subunits, POMP, PAC1 (a monitor of the PAC1/PAC2 heterodimer), PAC3 (a monitor of the PAC3/PAC4 heterodimer), and PI31. Consistent with results from the activity and western blotting assays described above, depletion of POMP significantly reduced the content of intact proteasome holoenzymes in both PI31 wild-type and knockout cells regardless of interferon-γ treatment (Figure 6C, lanes 5-8, and 13-16). This effect likely reflects a common role for POMP in the early-stage assembly of both constitutive and immuno-20S proteasomes. As expected, the content of Band 2, the late-stage POMP-containing assembly intermediate that accumulates in IFNγ-treated PI31 KO cells, was also significantly reduced in POMP knock-down samples (Figure 6C, lanes 21-24). In place of Band 2, POMP knock-down samples featured a prominent faster migrating band (annotated Band 3) containing 20S α subunits, PAC1 and PAC3 (Figure 6C, lanes 5-8, 29-32, and 37-40). The presence and composition of Band 3 in the absence of POMP indicate that Band 3 represents the α-subunit ring assembly intermediate that forms prior to the action of POMP and incorporation of β subunits. Accordingly, a PAC3-containing band that migrated more rapidly than Band 3, was detected in extracts from IFNγ-treated PI31 KO cells with normal POMP content (Figure 6C, lanes 33-36). This finding is consistent with current models in which PAC3/PAC4 ejection from the α-ring-POMP-PAC1/PAC2 complex is coincident with initial insertion of βi subunits on the assembled α ring (Figure 6C, lanes 33-36, annotated Band 4). Because PI31 had no effect on the content of Band 3 in POMP-depleted cells, its role in immuno-proteasome assembly is likely exerted at a step after the PAC1/PAC2-PAC3/PAC4-mediated formation of the α subunit ring. Such steps might include insertion of one or more βi subunits into the progressively growing assembly complex, or the joining of two 15S half-proteasomes. Thus, the general loss of β subunit content in POMP knock-down cells and the enhanced loss of βi subunit content upon depletion of both POMP and PI31 may reflect the instability of these subunits when their insertion into assembly complexes is blocked. In fact, a band likely containing β subunits (annotated Band 5 in Figure 6C, lanes 13 and 14) was present at much lower levels in when POMP was depleted from PI31 KO cells than from PI31 wild type cells (Figure 6C, lanes 9-16, see Discussion). Unlike POMP, PAC1, and PAC3, we were unable to detect PI31 as a component of component of Band 2, Band 3, or any other identifiable assembly intermediate. PI31 migrated near the dye front, a position that was indistinguishable from that of isolated, purified PI31 (Figure 6C, lanes 41-42 and 45-46). Similarly, PI31 was not detectably associated with assembly intermediates identified by glycerol density gradient centrifugation (Figure 5). These findings suggest that any role of PI31 in the assembly of IFNγ-induced immuno-proteasomes is characterized by highly transient interactions with intermediate complexes (see Discussion).

## DISCUSSION

The formation of large multisubunit proteins often requires chaperones to coordinate the spatial and temporal complexities of the assembly process. The tightly orchestrated assembly of the 28-subunit eukaryotic 20S proteasome is aided by five dedicated chaperones: POMP and two heterodimeric proteins, PAC1/PAC2 and PAC3/PAC4 ^40,60–63^. Moreover, N-terminal pro-peptides of several constituent catalytic β subunits contribute additional chaperone-like functions to 20S assembly^40^. Extensive genetic, biochemical, and structural studies have provided a detailed and increasingly comprehensive understanding of the pathway, and mechanisms, and relative roles of each protein in the assembly of the 20S proteasome (Figure 7).

**Figure 7.**
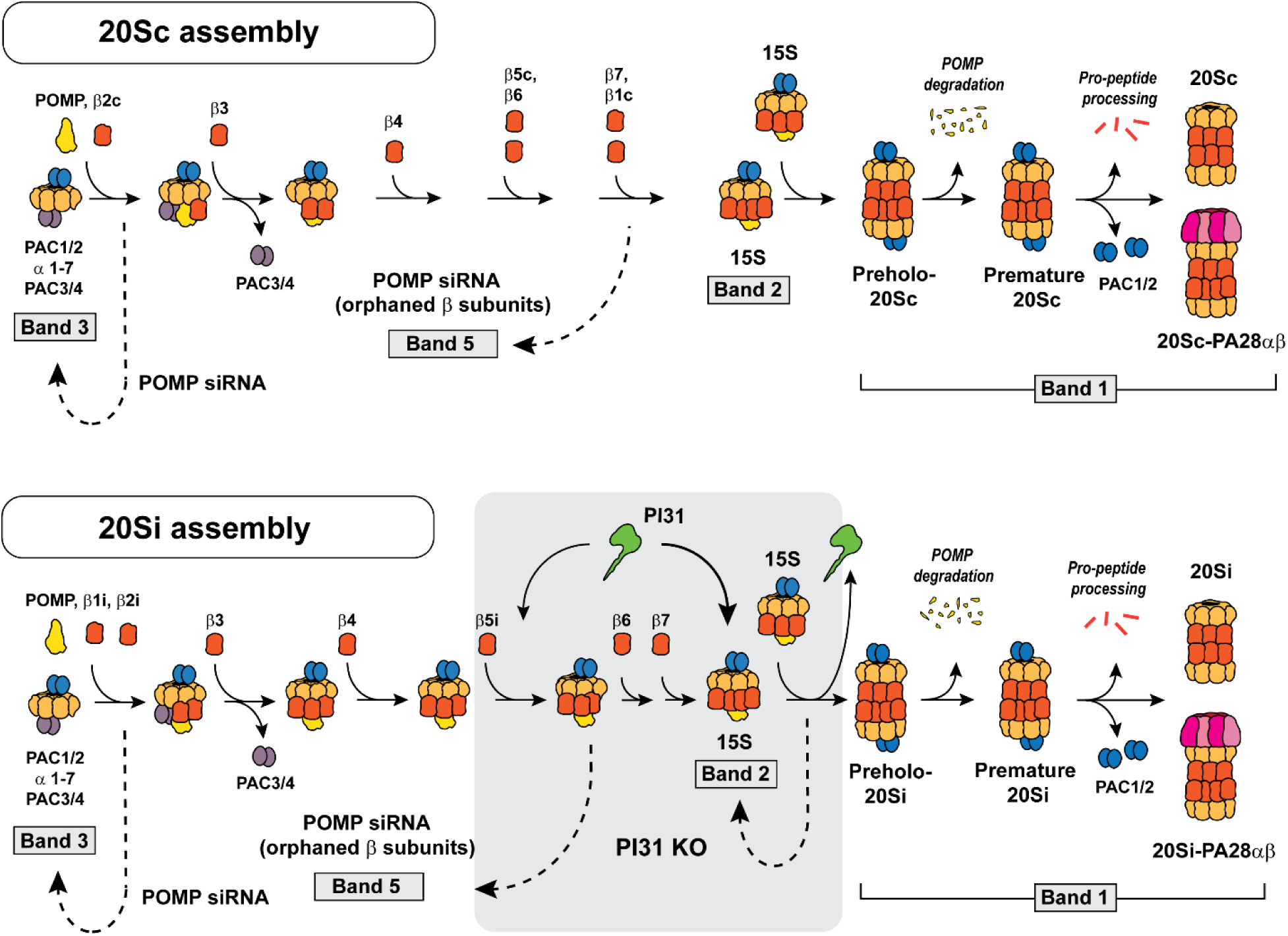
Models of 20Sc and 20Si proteasome assembly. **(Upper),** Model of 20Sc assembly consistent with current literature. **(Lower),** Model of IFNγ-induced 20Si assembly. Gray box indicates possible sites of PI31 action based on the present study. Denoted bands correspond to complexes identified on native gels (Figures 4 and 6) and discussed in the text.

Higher eukaryotes can express multiple 20S proteasome isoforms characterized by unique complements of genetically and functionally distinct catalytic β subunits. Although the assembly pathways of these different 20S isoforms are broadly similar, specific aspects of the process appear to differ. Such distinctions may be the basis for the selective formation of given isoforms under different physiologic conditions. For example, the selective formation of immuno-proteasomes upon interferon-γ-induced expression of βi subunits seems to be driven by the initial preferential addition of β1i and β2i to the α subunit ring in lieu of β2c^64^ This process appears to divert subsequent steps of 20S assembly in favor of 20Si (Figure 7).

The current work identifies PI31, a poorly understood proteasome-binding protein, as a positive regulator of 20Si assembly. Depletion of cellular PI31 by genetic knockout or knockdown methodologies stalls interferon-γ-induced 20Si assembly at an intermediate stage similar or identical to the well-characterized 15S half-proteasome assembly complex (Figure 4, Band 2). This finding suggests that PI31 plays a role in a process required for late-stage insertion of one or more β subunits into the complex and/or for fusion of two fully-assembled half-proteasomes to form the pre-holo 20Si complex (Figure 7). The reduced content of mature 20Si proteasomes caused by the absence of PI31 limits the formation of functional 20Si-PA28 and 26Si proteasome holoenzymes. Notably, the majority of reduction in IFNγ-induced immuno-proteasome activity in PI31 KO cells is accounted for by decreasing levels of 20Si-PA28 holoenzymes. Although the preferential formation of this holoenzyme in wild-type cells may be driven, in part, by the IFNγ-increased gene expression of PA28α and PA28β subunits ^65^, PA28 levels do not appear to be a limiting factor for holoenzyme formation under any condition examined here. It is unclear whether some aspect of interferon-γ signaling normally promotes 20Si-PA28 formation or whether any factor other than lower 20Si levels *per se* contributes to reduced 20Si-PA28 levels in PI31 KO cells.

Despite the robust effect of PI31 knockout on decreasing cellular immuno-proteasome content and activity, neither the mechanism of this effect nor the basis for its apparent specificity for the immuno-proteasome is clear. Unlike established 20S assembly chaperones POMP, PAC1/2, and PAC3/4, PI31 was not a detectable component of any assembly complex resolved here. This may be a consequence of transient or weak interactions of PI31 with these complexes that preclude their stable capture by native PAGE or density gradient centrifugation methodologies. Moreover, like the degradation of POMP that occurs upon maturation of 20S, the selective degradation of PI31 by 20Si, such as demonstrated by *in vitro* studies ^58^, may be a normal element of the mechanism of 20Si assembly and contribute to our failure to detect PI31 in assembly complexes. Regardless of the inability to detect a physical association of PI31 with assembly intermediates, several structural and functional properties of PI31 lead us to posit that PI31 normally functions as a chaperone for 20Si assembly via direct interactions with proteasome subunits. First, high-resolution cryo-EM structures of a 20S-PI31 complexes reveal extensive interactions of PI31 with 20S proteasome subunits. These interactions include those between PI31’s carboxyl terminus and each catalytic β subunit ^46–48^. Although these interactions may be unique to fully assembled 20S proteasomes, it is reasonable to assume that the same, or closely related interactions also occur between PI31 and proteasome assembly complexes or individual subunits. Such interactions may result in the recruitment or delivery of one or more β subunits to assembly complexes and/or in induction of conformational changes within assembly complexes required for progression of the assembly process. Interactions between PI31 and individual β subunits, as might be involved in a subunit delivery to an assembly intermediate, would likely not be detected at the resolution of native PAGE or glycerol density gradient centrifugation. In such a case, the absence of PI31 might promote turnover of the unincorporated subunit that accounts for its diminished content by western blotting. Second, we recently identified differential interactions of PI31 with the catalytic β subunits of intact 20Sc and 20Si proteasomes ^58^. These differences provide a potential basis for the selectivity of PI31 on 20Si assembly. For example, specific features of PI31 to βi subunits may direct their selective incorporation into assembly complexes. Alternatively, preferential binding of PI31 to βc subunits may sequester βc subunits from productive association with assembly complexes, thereby favoring incorporation of βi subunits into newly synthesized proteasomes. Such an inhibitory effect of PI31 on formation of 20Sc, however, would likely demand additional forms of regulation to prevent interference of PI31 with normal assembly of 20Sc in the absence of interferon-γ. Third, PI31 harbors an HbYX motif at its carboxyl terminus. This motif is found in other 20S regulatory proteins where it supports holoenzyme stability and induces opening of the substrate access pore ^66,67^. HbYX motifs, however, are also found in PAC1 and in yeast Pba1/Pba2 20S chaperones where they play specific roles in 20S assembly^40^. Because the HbYX motif is not required for PI31’s binding to or inhibition of intact 20S proteasomes, its conservation in PI31 may indicate a role analogous to that of other assembly chaperones^50^. Finally, we note a previous report that demonstrated inhibition of IFNγ-induced immuno-proteasome activity after massive overexpression of PI31 ^51^. Similar to the current results for PI31 knockout, this effect was accompanied by reduced assembly of immuno-proteasomes and an accumulation of unprocessed pro-peptide forms of βi subunits in assembly intermediates ^51^. Although no mechanism for this effect was described, similar phenotypes are often caused by both depletion or overexpression of proteins that function as scaffolds and coordinate interactions among different proteins. We have also observed inhibition of IFNγ-induced proteasome activity upon massive PI31 overexpression in several cell lines (Supplemental Figure 6). This inhibition was not accompanied in our studies by a detectable accumulation of unprocessed βi subunits. Moreover, unlike PI31 knockout, PI31 overexpression also inhibited constitutive proteasome activity in both interferon-γ treated and untreated cells. Thus, the mechanism(s) by which PI31 overexpression inhibits proteasome activity under these conditions remains unclear.

Although we favor a model in which PI31 supports 20Si assembly via direct interactions with proteasome subunits or chaperones, we cannot exclude models based on other PI31 properties and functions. For example, PI31 is known to bind to FbxO7, a subunit of an SCF-type E3 ligase ^68^. Thus, an FbxO7-PI31-containing ligase can participate in the targeting substrates for ubiquitin-dependent proteolysis ^69^. A comprehensive list of client substrates targeted by this ligase is unavailable, but hypothetically might include a protein whose degradation is required for normal 20Si assembly. FbxO7 also has been reported to have E3 ligase-independent functions, but possible roles of PI31 in these roles are unclear ^69^. We also cannot exclude the possibility that effects of PI31 on 20Si assembly reported here are linked to features of IFNγ-stimulated proteasome biogenesis distinct from the induced transcription and synthesis of βi subunits. For example, PI31 may regulate or be regulated by a component or activity of the interferon-γ signaling pathway that licenses its role in 20S assembly. Likewise, PI31 may be required to support enhanced rates of proteasome assembly such as those that may occur in response to interferon-γ rather than to promote the assembly of a specific proteasome isoform. These speculations and other possible determinants of PI31action on proteasome assembly will require additional investigation.

The current results add another function to the growing list of reported physiologic roles for PI31. In addition to its original description as an *in vitro* inhibitor of purified 20S proteasome, PI31 has been described as a direct activator or 26S proteasome^52^, an assembly factor for 26S proteasomes ^55^, a negative regulator of antigen presentation ^51^, an adaptor for dynein-mediated proteasome transport on neuronal axons ^56^, a negative regulator of viral replication ^53^, and a regulator of mitochondrial function ^70^. We propose that the unique structure of PI31 may enable it to participate in multiple and diverse processes whose given manifestations depend on specific cellular and physiologic conditions.

## MATERIALS AND METHODS

### Generation of PI31 knockout cell lines

The PSMF1 gene (UniProt ID Q92530) encoding PI31 was disrupted in HAP1 and HepG2 cells with sgRNA 5’-CCTTGTGAAAGCCATCACCG-3’, and PAM sequence TGG (GenScript) for HepG2 cells and sgRNA 5’-CATCCTTATACTCATACCG-3’ for HAP1 cells, each targeting exon 2. Cells were plated into 6-well plates and transfected with 5 µl Lipofectamine 3000 and 5 µl P3000 (Invitrogen) containing 1 µg pLentCRISPR v2 plasmid. Twenty-four hours after transfection, cells were exposed to selection media with 5 μg/l puromycin for an additional 48 hours. Selection media was removed, and monoclonal cells were screened by limiting dilution. Gene disruption was confirmed by Sanger sequencing and western blotting. Sequencing data demonstrate a 10 base pair deletion for HAP1 cells and a 1 base pair deletion for HepG2 cells in exon 2 of the PSMF1 gene (Supplemental Figure 1).

### Cell culture

Human cell lines (HAP1, HepG2, and HeLa) were obtained from American Type Culture Collection. Cells were cultured in IMDM (Hap1) and DMEM (HepG2 and HeLa) media supplemented with glutamine, 10% fetal bovine serum, and penicillin (50 IU/ml)/streptomycin (50 μg/ml), at 37°C with 5% CO_2_.

### Interferon-γ treatments

Cells were treated with 100 U/ml recombinant human interferon-γ (Thermo-Fisher) as indicated in specific experiments. Prior to treatments, cells were plated at densities of 1 – 1.5 × 10^6^ cells per 100 mm plate and exposed the following day to for durations indicated in the figure legends.

### Preparation of cell extracts

Cell extracts were prepared for proteasome assays, gel electrophoresis, and glycerol density gradient centrifugation. Cells were washed three times with ice-cold PBS. Washed cells were collected in hypotonic lysis buffer (20 mM Tris-HCl (pH 7.6 at 37°C), 20 mM NaCl, 1 mM DTT, 5 mM MgCl_2_, 100 µM ATP, 0.1% NP-40). For analyses of proteasome assembly intermediates in experiments where activity was not measured, 10 µM MG132 was added to the lysis buffer. Cells were mechanically lysed by 10-15 passes through a 27½ G needle, and supernatants were collected following centrifugation at 16,000 rcf for 20 minutes at 4°C. Samples subjected to native PAGE were centrifuged at 20,000 rpm for 20 minutes, 4°C. Protein content of cell lysates was determined using detergent-compatible BCA assays (Pierce BCA Protein Assay Kit, ThermoScientific) as described by the manufacturer. Protein concentrations were determined using a standard curve of diluted BSA standards.

### Measurement of proteasome activity using fluorogenic peptide substrates

Proteasome activity was measured in cell extracts by monitoring the fluorescence of 7-amino-4-methylcourarin (amc) cleaved from amc-linked peptide substrates Suc-LLVY-amc, Ac-ANW-amc, or Ac-PAL-amc. 10 μl of cell extract were added to 150 µl of 50 µM peptide substrate in an assay buffer consisting of 20 mM Tris-HCl, pH 7.6, 20 mM NaCl, 5 MgCl_2_ and 100 mg ATP. Assays were conducted in triplicate in 96 well plates using a BioTek Synergy plate reader at 37°C. Fluorescence (excitation 380 nm, emission 460 nm) was measured every 45 seconds for 21 minutes. Control assays contained lysis buffer without cell extract. Rates of production of amc fluorescence were determined by instrument software and expressed as arbitrary fluorescent units (AFU)/min/μg extract proteins. All reported rates are from assays in which amc production was linear with respect to incubation time and with respect to extract concentration.

### Measurement of proteasome activity using proteasome activity probes

Proteasome activity was also measured using activity-based probe labeling of Me_4_BodipyFL-AHX_3_Leu_3_VS for each active site. Whole cell lysates were incubated in 1 µM activity-based probe at 37°C for one hour. Reactions were terminated by the addition of 5x SDS sample buffer. Extracts were subjected to SDS-PAGE, transferred to nitrocellulose membranes, and imaged on an Odyssey M imager (LICOR) at 520 nm.

### Western Blotting

Cell extracts were denatured in SDS sample buffer under reducing conditions and denatured at 100 °C for 5 minutes. Known amounts of protein were separated by SDS-PAGE (10-12% acrylamide) and transferred to 0.2 µm nitrocellulose membranes in transfer buffer containing 20% MeOH. Membranes were then blocked in 5% milk in TBST buffer for 1 hour at room temperature. Primary antibodies were incubated overnight at 4°C. Secondary antibodies with IRDye were obtained from LiCOR and used at 1:10,000 dilution. Near-infrared emission from IRDye was imaged using Odyssey Dlx or Odyssey M (LICOR) at channel 700 or 800 nm depending on the secondary antibody.

The following primary antibodies were used at 1:2,000 to 1:500 dilutions:

Commercial antibodies
20Sc β2 subunit (Rabbit monoclonal IgG; Cell Signaling Technology #13207)
20Sc β2i subunit (Rabbit polyclonal; Cell Signaling Technology #78385)
20S α4 subunit (Mouse monoclonal IgG1, MCP34; Enzo BML-PW8120),
20S α1-3/5-7 (Mouse monoclonal IgG1 (MCP231); Abcam ab22674),
PI31 (Rabbit polyclonal; Abclonal BML-PW9710),
PAC1 (Rabbit polyclonal; Cell Signaling Technology #13378),
PAC3 (Mouse IgG2b; Proteintech #67466-1-Ig)
POMP (Rabbit monoclonal IgG; Cell Signaling Technology #15141),
Actin (mouse monoclonal C4; Sigma-Aldrich MAB1501),
β-Tubulin (Rabbit polyclonal; Cell signaling Technology #2146),
Flag (mouse monoclonal; ThermoFisher Scientific #MA1-91878),
Ubiquitin (Mouse monoclonal (FK2); Sigma-Aldrich #ST1200).
20S α2 subunit (Mouse monoclonal IgG MCP21, was gift from Klaus Hendil)

Polyclonal antibodies against human proteins generated in rabbits were prepared in the DeMartino laboratory against indicated antigens.

1. β1 subunit (antigen peptide: N-LAAIAESGVERQVLLGDQIPKFAVATLPPA-C)
2. 20S, β5 subunit (antigen peptide: N-WIRVSSDNVADLHEKYSGSTP-C)
3. 20S, β1i subunit (antigen peptide: N-IYLVTITAAGVDHRVILGNELPKFYDE-C
4. β5i subunit (antigen peptide: N-WVKVESTDVSDLLHQYREANQ-C
5. E446 protein (antigen purified 20S proteasome)^71^
6. PA28αβ protein (antigen purified PA28αβ)^72^
7. Rpt2 subunit (antigen peptide: N-ENVLYKKQEGTPEGLYL-C)
8. Rpt5 subunit (antigen peptide: N-ILEVQAKKKANLQYYA-C)

### Native polyacrylamide gel electrophoresis (PAGE)

Native polyacrylamide gel electrophoresis was conducted using 3-8% polyacrylamide Tris-acetate gels (Life Technologies) as described previously ^73,74^. In brief, samples were mixed with 5x native PAGE sample buffer, applied to the loading wells and electrophoresed for 2.5 – 3.0 hours at 100 V at 4°C with a 90 mM Tris-borate running buffer containing 5 mM MgCl_2_ and 100 μM ATP. Following electrophoresis, proteasome activity of separated bands was performed by incubating gels in 50 µM fluorogenic peptide substrate (Suc-LLVY-amc, Ac-ANW-amc, or Ac-PAL-amc in the presence of 5 MgCl_2_ and 100 μM ATP. Incubations were conducted at 37°C for 30 minutes. Gels were imaged using a ChemiDoc UV transilluminator (BioRad) using the fluorescein channel. For immunoblotting of native PAGE samples, gels were transferred and blotted under the same conditions described for western blotting above.

### Second dimension SDS polyacrylamide gel electrophoresis

Second dimension SDS PAGE was conducted by excising individual lanes of native polyacrylamide gels. Excised gel lanes were incubated in 1x SDS sample buffer at room temperature for 10 minutes and adhered horizontally to the top of SDS 12% polyacrylamide gel with electrode buffer containing 0.5% agarose. After transfer, first dimension native lane boundaries were visualized by Ponceau S staining and marked prior to blocking. SDS-PAGE and western blotting were conducted as described above.

### Glycerol Density Gradient Centrifugation

Glycerol density gradient centrifugation was conducted, as described previously ^75^. Cell extracts (approximately 500 μg total extract protein) were applied to a 1.95 ml 10-40% linear glycerol gradient in a buffer containing 50 mM Tris-HCl (pH 7.6, 4°C), 100 μM ATP, 5 mM MgCl_2_, 1 mM DTT. Centrifugation was carried out for 3.5 hrs at 4°C in a Beckman Optima TL ultracentrifuge at 55,000 rpm in TLS-55 rotor. 21 fractions (100 μl/fraction) were collected and analyzed as described for given experiments.

### RNA interference

RNA interference (RNAi) for knockown of POMP and PI31 were conducted using ON-TARGETplus SMARTpool siRNA mixes (Horizon Discovery; catalog #L-016844-01-0005 and #L-012320-01-0005 respectively). Pooled siRNA mixes contained four individual target sequences which were also validated separately. Specific siRNAs used individually for POMP were: GGGUCUAUUUGCUCCGCUA and UCAUGAUCUUCUUCGGAAA. Specific siRNAs for PI31 were: CAACAUAUAUCCUCGACCA and GAGUGGAAAGAGCACGAUA. 25 nM siRNAs were transfected into cells using DharmaFECT1 according to manufacturer’s recommendations for 6-well plate scale. A pool of control non-targeting siRNAs was used as a negative control (Horizon Discovery; #D-001810-10-05). PI31 and POMP protein expression was assessed at 24 and 48 hours to determine an appropriate time course. Following PI31 or POMP knockdown, cells were exposed to control or human interferon-γ (100 U ml^-1^) for 24 hours. Cells were harvested and assayed as described above.

### Overexpression of PI31

Overexpression of PI31 was performed as described previously (Hsu *et al*., 2023). Briefly, cells were transfected with Lipofectamine 3000/P3000 (Invitrogen) with a pCMV3-N-FLAG-PI31 construct (HG17079-NF, Sino Biological Inc.). 24 hours after transfection, transfection media was removed, and cells were exposed to IFNγ or control treatments for the amounts of time indicated in figure legends.

### Statistics

Experiments were repeated independently for the number of biologic replicates indicated in figure legends. Treatment effects were analyzed by repeated measures ANOVA matched within experiments. Where significant treatment effects were found, Tukey’s HSD post-hoc test was performed to determine pairwise differences. All statistics were calculated using GraphpadPrism Software (v10.2.3).

## ACKNOWLEDGEMENTS

This work was supported by a grant from the National Institutes of Health (GM129088) to G.N.D.

