## Supplemental Figures for "PI31 is a positive regulator of 20S immunoproteasome assembly"

Department of Physiology

University of Texas Southwestern Medical Center

Dallas, TX  75390-9040

**A.**

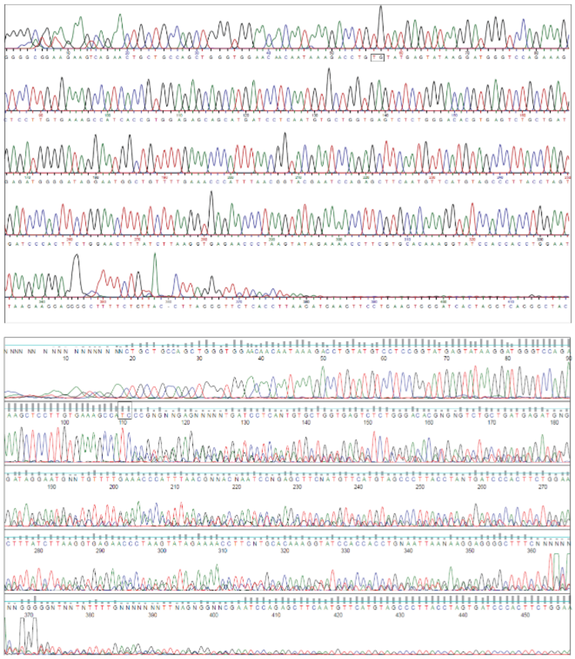


**B.**

```
                                  << Exon 2 >>                            |----gRNA (HAP1)----|
NC_000020.11:1125529-1125804      TGCCAGCTGGGTGGAACAACAATAAAGACCTGTATGTCCTCCGGTATGAGTATAAGGATG      60
HepG2-WT                          TGCCAGCTGGGTGGAACAACAATAAAGACCTGTATGTCCTCCGGTATGAGTATAAGGATG      60
HepG2-KO                          TGCCAGCTGGGTGGAACAACAATAAAGACCTGTATGTCCTCCGGTATGAGTATAAGGATG      60
HAP1-KO                           TGCCAGCTGGGTGGAACAACAATAAAGACCTGT----------GTATGAGTATAAGGATG      50
                                  *********************************          *****************

                                               |---gRNA (HepG2)----|
NC_000020.11:1125529-1125804      GGTCCAGAAAGCTCCTTGTGAAAGCCATCACCGTGGAGAGCAGCATGATCCTCAATGTGC      120
HepG2-WT                          GGTCCAGAAAGCTCCTTGTGAAAGCCATCACCGTGGAGAGCAGCATGATCCTCAATGTGC      120
HepG2-KO                          GGTCCAGAAAGCTCCTTGTGAAAGCCATC-CCGTGGAGAGCAGCATGATCCTCAATGTGC      119
HAP1-KO                           GGTCCAGAAAGCTCCTTGTGAAAGCCATCACCGTGGAGAGCAGCATGATCCTCAATGTGC      110
                                  ***************************** ******************************

                                   |Intron >>
NC_000020.11:1125529-1125804      TGGTGAGTCTCTGGGACACGTGAGTCTGCTGATGAGATGGGGATAGGAATGGCTGTTTTG      180
HepG2-WT                          TGGTGAGTCTCTGGGACACGTGAGTCTGCTGATGAGATGGGGATAGGAATGGCTGTTTTG      180
HepG2-KO                          TGGTGAGTCTCTGGGACACGTGAGTCTGCTGATGAGATGGGGATAGGAATGGCTGTTTTG      179
HAP1-KO                           TGGTGAGTCTCTGGGACACGTGAGTCTGCTGATGAGATGGGGATAGGAATGGCTGTTTTG      170
                                  ************************************************************

NC_000020.11:1125529-1125804      AAACCCATTTAACGGTACGAATCCAGAGCTTCAATGTTCATGTAGCCCTTACCTAGTGAT      240
HepG2-WT                          AAACCCATTTAACGGTACGAATCCAGAGCTTCAATGTTCATGTAGCCCTTACCTAGTGAT      240
HepG2-KO                          AAACCCATTTAACGGTACGAATCCAGAGCTTCAATGTTCATGTAGCCCTTACCTAGTGAT      239
HAP1-KO                           AAACCCATTTAACGGTACGAATCCAGAGCTTCAATGTTCATGTAGCCCTTACCTAGTGAT      230
                                  ************************************************************

NC_000020.11:1125529-1125804      CCCACTTCTGGAACTTTATCTTAAGGTGAGAACCCT      276
HepG2-WT                          CCCACTTCTGGAACTTTATCTTAAGGTGAGAACCCT      276
HepG2-KO                          CCCACTTCTGGAACTTTATCTTAAGGTGAGAACCCT      275
HAP1-KO                           CCCACTTCTGGAACTTTCTCTTAAGGTGAGAACCCT      266
                                  ***************** ******************
```

**Supplemental Figure 1. Validation of PI31 knockout from HAP1 and HepG2 cells.** The PSMF1 encoding PI31 was disrupted as described in Materials and Methods. **(Panel A),** Sanger sequencing results for PI31 knockout lines. (Upper), PI31-KO in HAP1 cells resulted from a 10 bp deletion between the indicated T and G nucleotides. (Lower), PI31-KO in HepG2 cells resulted from a 1 bp deletion after the indicated TC. Deletions are indicated by boxes. **(Panel B),** DNA sequences were aligned with the reference genome and the independently-sequenced HepG2 wildtype sequence using Clustal Omega online tool (https://www.ebi.ac.uk/jdispatcher/msa/clustalo). The boundary between the PSMF1 exon 2 and intron region is shown. Guide RNAs used to knockout PI31 from respective cell lines are underlined, and deleted sequences are highlighted in yellow.

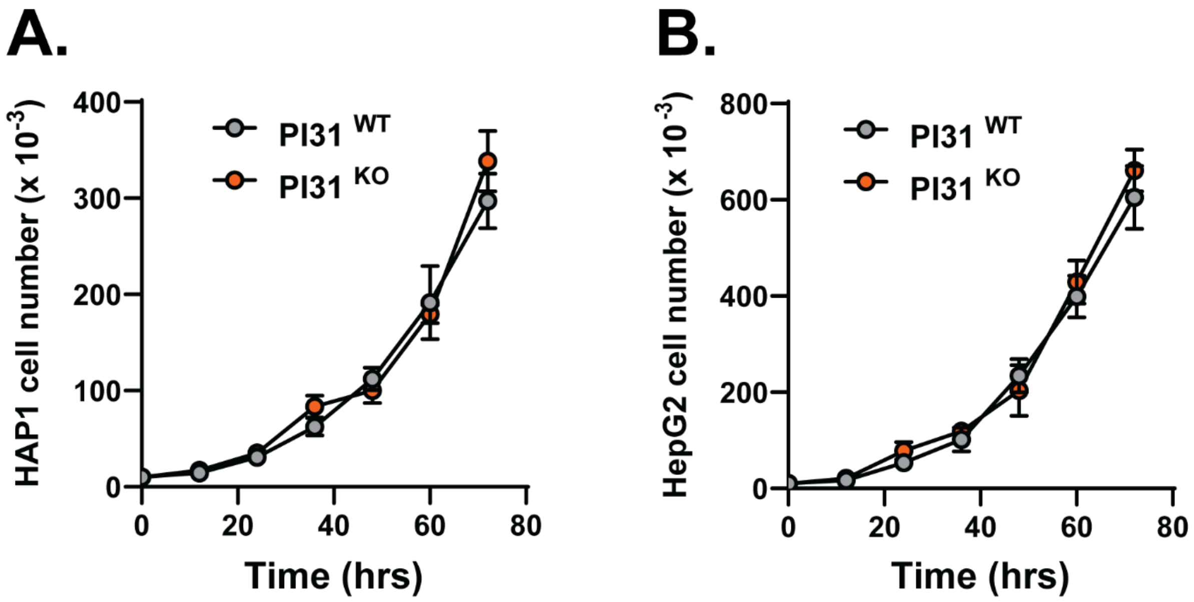


**Supplemental Figure 2. PI31 knockout does not affect cell proliferation.** PI31 wild-type and knockout HAP1 **(Panel A)** and HepG2 **(Panel B)** cells were seeded at 10,000 cells per plate and incubated under standard culture conditions. Culture media was changed every 24 hrs. At indicated times, cells were harvested and counted. Data points represent mean cell number $\pm$ s.d. of triplicate plates.

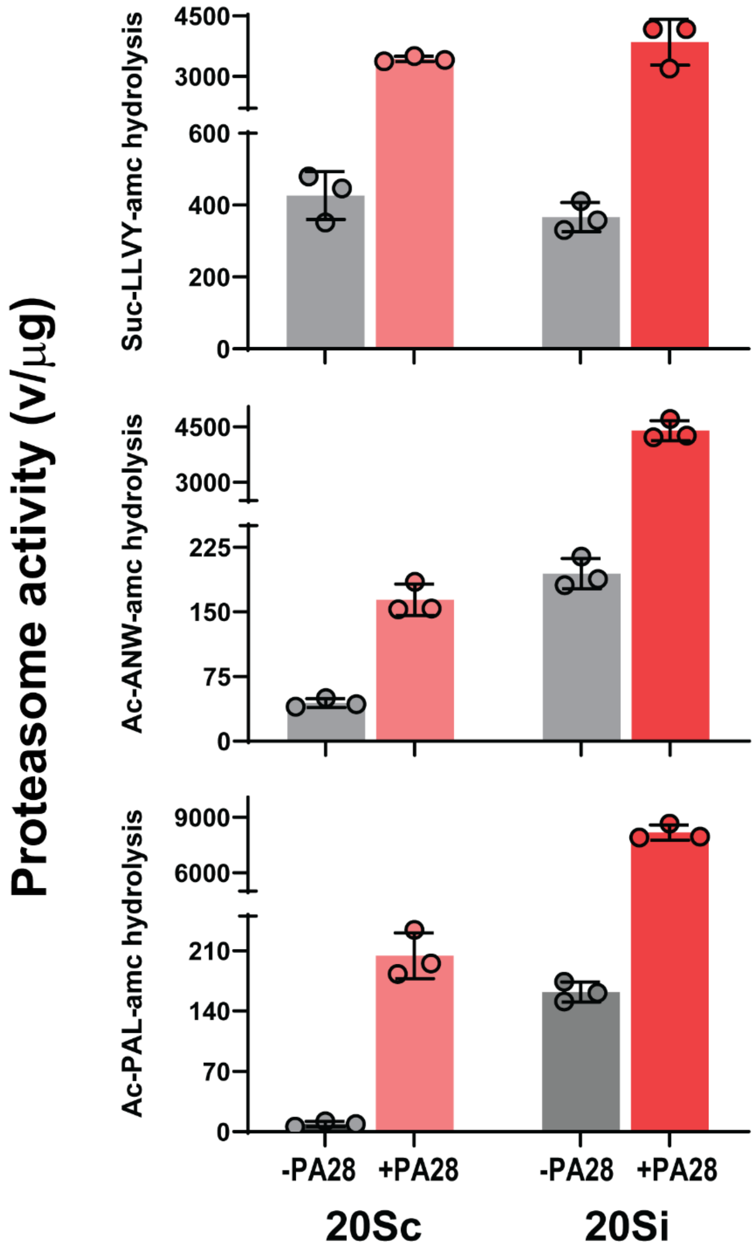


**Supplemental Figure 3. Relative substrate specificities of constitutive and immuno- 20S proteasomes.** Constitutive (20Sc) and immuno- 20S (20Si) proteasomes were purified from bovine red blood cells or bovine spleen, respectively, and assayed for hydrolysis of indicated peptides in the presence or absence of purified PA28αβ, as described previously[58]. Bars represent mean rates of hydrolysis $\pm$ s.d. of triplicate assays. Similar results were obtained with two independent preparations of purified proteasomes.

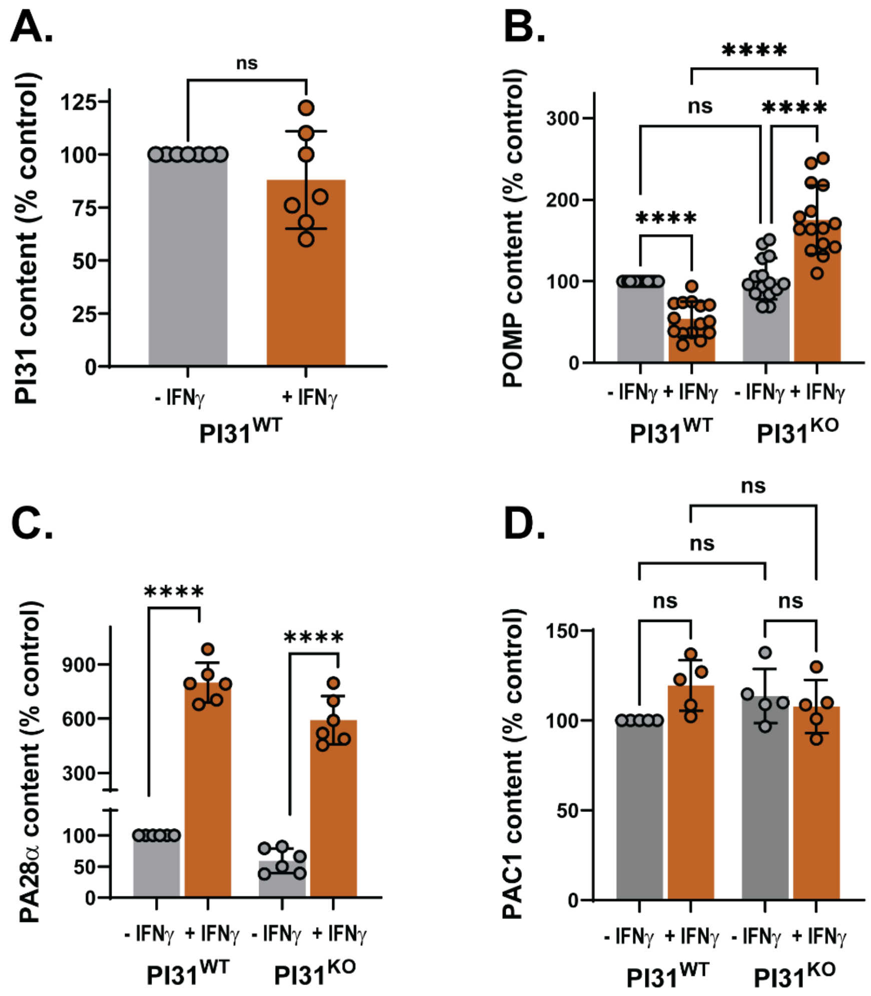


**Supplemental Figure 4. Knockdown of PI31 by RNAi inhibits interferon-γ induced increases of immuno-proteasome content and activity in HepG2 cells.** HepG2 cells were transfected with a non-coding siRNA (NC), an siRNA targeting PI31 (PI31), or transfection reagent only (-). After 24 hrs, cells were treated with 100 U/ml of interferon-γ for 48 hrs. Cells extracts were prepared as described under Materials and Methods. (**Panel A),** Cell extracts for indicated conditions were normalized for total protein and subjected to western blotting for indicated proteins. Lanes show blots from two independent biologic experiments. **(Panel B**), Extracts, normalized for total protein, were assayed for proteasome activity using the indicated peptide substrates. Rates of substrate hydrolysis for cells treated with non-coding (NC) siRNA were set to 100 and rates for all other treatments were expressed relative to that value. Each data point represents the mean value of triplicate assays for a given biologic experiment. Bars represent mean values $\pm$ s.d. of independent biologic experiments. Differences were analyzed by repeated measures 2-way ANOVA and Tukey's HSD posthoc test (* $p<0.05$; ** $p<0.01$; *** $p<0.001$). **(Panel C)**, Extracts of cells from indicated conditions were normalized for total protein and subjected to glycerol density gradient centrifugation as described in Materials and Methods. Gradient fractions were assayed for immuno-proteasome activity using Ac-ANW-amc substrate. 20S-PA28 and 26S show sedimentation positions of respective purified holoenzyme standards.

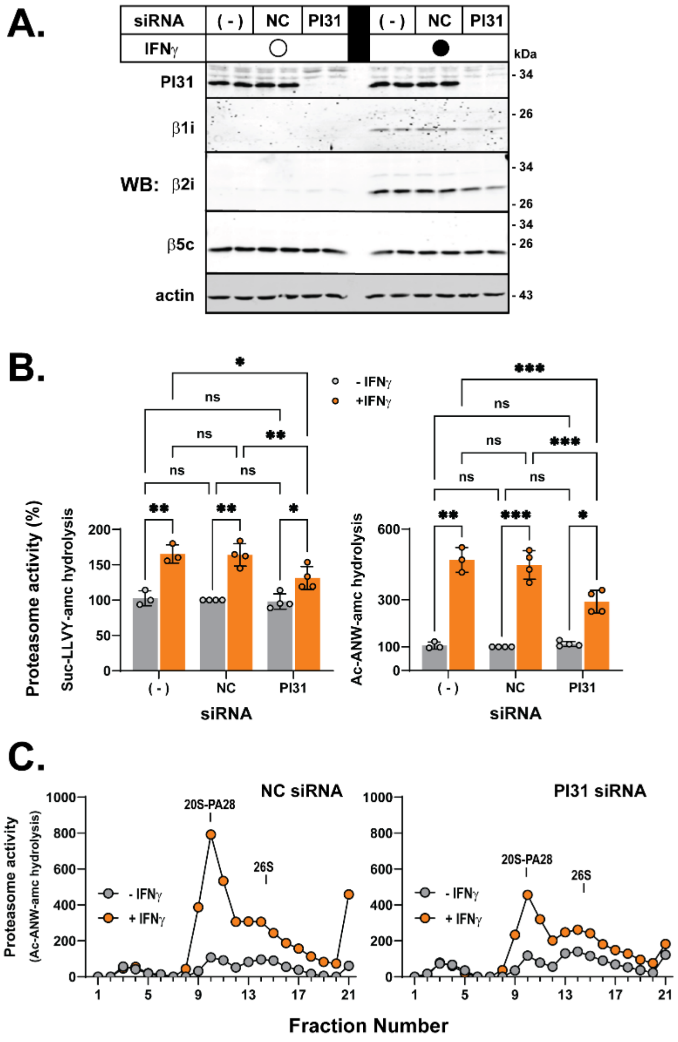


**Supplemental Figure 5. Effect of PI31 knockout on regulatory proteins of the proteasome.** PI31 wild-type and PI31 KO HAP1 cells were cultured in the absence (-IFNγ) or presence (+IFNγ) of 100 U/ml human interferon-γ for 24 hrs as indicated. Extracts were prepared as described in Materials and Methods, normalized for total protein and subjected to western blotting for indicated proteins. Blots for PI31 **(Panel A)**, POMP **(Panel B)**, PA28α **(Panel C)**, and PAC1 **(Panel D)** were quantified using ImageStudio (LiCOR) software. Within each independent experiment, the protein level from PI31 WT cells in the absence of interferon-γ was set at a value of 100 and all other values were expressed relative to that. Individual data points represent independent biologic experiments. Differences were analyzed by repeated measures 1- or 2-way ANOVA and Tukey's HSD posthoc test where appropriate (*ns* p>0.05; **** p<0.0001).

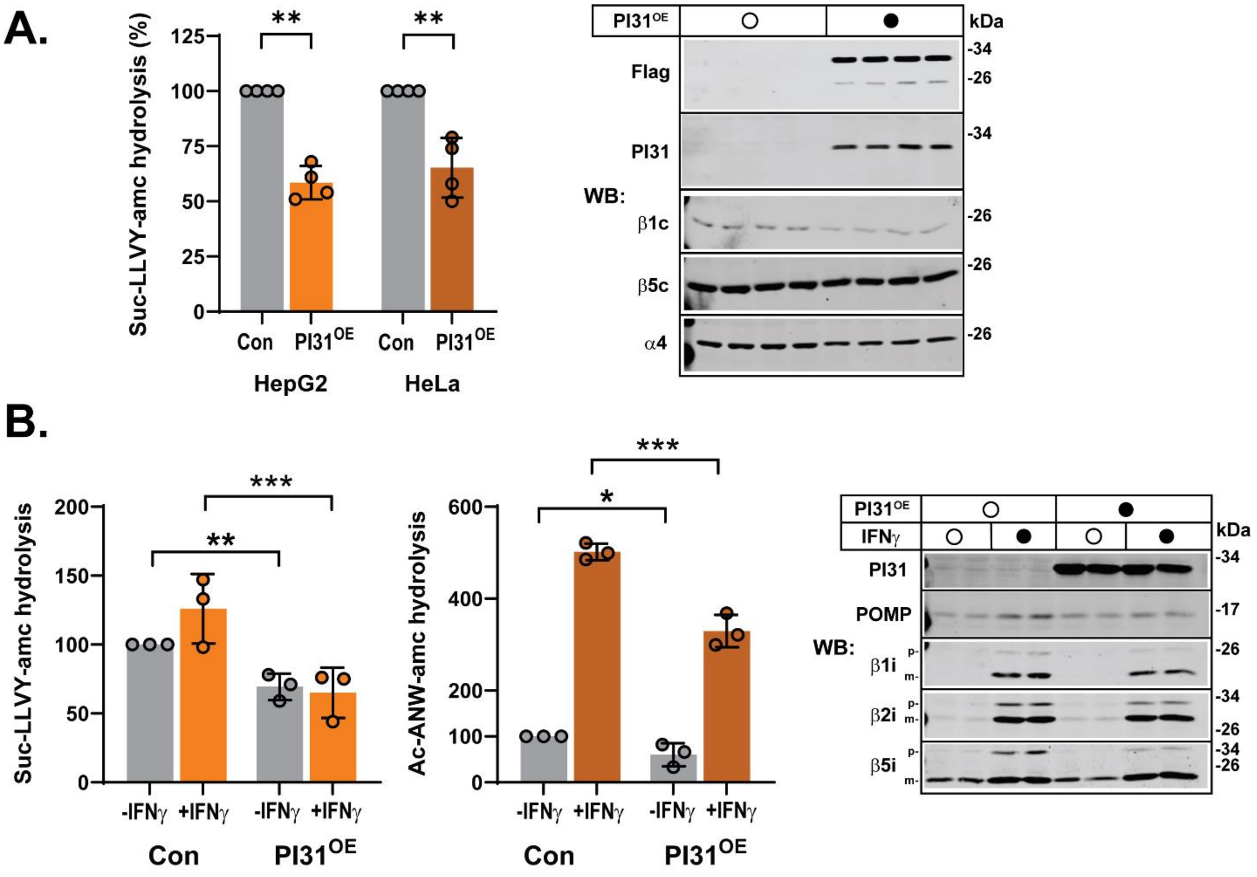


**Supplemental Figure 6. Massive overexpression of PI31 inhibits constitutive and IFNγ-induced immuno-proteasome activities in HepG2 and HeLa cells.** HepG2 and HeLa cells were transfected with Flag-PI31 ($PI31^{OE}$) or transfection reagent (Con) as described in Materials and Methods. **(Panel A),** 48 hrs after transfection extracts of HepG2 or HeLa cells were normalized for total protein and assayed for constitutive proteasome activity (left) or subjected to western blotting for indicated proteins (right). Bars represent mean values ± s.d. of indicated independent experiments. Control values were assigned values of 100 and PI31 overexpression samples were expressed relative to that. Differences were analyzed by repeated measures ANOVA and Tukey's HSD posthoc test ** $p<0.01$. Western blots show HeLa cell extracts from four independent experiments. **(Panel B),** HeLa cells were transfected with Flag-PI31 or vector alone. After 24 hrs, indicated cells were treated with 100 U/ml interferon-γ for an additional 24 hrs. Cell extracts were assayed for proteasome activity with the indicated substrates (left) or subjected to western blotting for the indicated proteins (right). Rates of substrate hydrolysis for cells treated with vector alone were set to 100 and rates for all other treatments were expressed relative to that value. Each data point represents the mean value of triplicate assays for a given biologic experiment. Bars represent mean values ± s.d. of independent biologic experiments. Differences were analyzed by repeated measures 2-way ANOVA and Tukey's HSD posthoc test (* $p<0.05$; ** $p<0.01$; *** $p<0.001$). Western blots show data from two independent biologic experiments. "p-" and "m" indicate positions of the unprocessed pro-peptide and mature forms, respectively, of indicated βi subunits.
